# Rational Design of Live Biotherapeutic Products for the Prevention of *Clostridioides difficile* Infection

**DOI:** 10.1101/2024.04.30.591969

**Authors:** Shanlin Ke, Javier A Villafuerte Gálvez, Zheng Sun, Yangchun Cao, Nira R Pollock, Xinhua Chen, Ciarán P Kelly, Yang-Yu Liu

## Abstract

*Clostridioides difficile* infection (CDI) is one of the leading causes of healthcare- and antibiotic-associated diarrhea. While fecal microbiota transplantation (FMT) has emerged as a promising therapy for recurrent CDI, its exact mechanisms of action and long-term safety are not fully understood. Defined consortia of clonal bacterial isolates, known as live biotherapeutic products (LBPs), have been proposed as an alternative therapeutic option. However, the rational design of LBPs remains challenging. Here, we employ a computational pipeline and three independent metagenomic datasets to systematically identify microbial strains that have the potential to inhibit CDI. We first constructed the CDI-related microbial genome catalog, comprising 3,741 non-redundant metagenome-assembled genomes (nrMAGs) at the strain level. We then identified multiple potential protective nrMAGs that can be candidates for the design of microbial consortia targeting CDI, including strains from *Dorea formicigenerans*, *Oscillibacter welbionis*, and *Faecalibacterium prausnitzii*. Importantly, some of these potential protective nrMAGs were found to play an important role in the success of FMT, and the majority of the top protective nrMAGs can be validated by various previously reported findings. Our results demonstrate a computational framework for the rational selection of microbial strains targeting CDI, paving the way for the computational design of microbial consortia against other enteric infections.

## Introduction

*Clostridioides difficile* infection (CDI) is one of the leading causes of healthcare- and antibiotic-associated diarrhea, affecting roughly 500,000 patients and leading to almost 30,000 deaths annually in the United States^1,2^. Exposure to toxinogenic *C. difficile* can lead to a spectrum of clinical outcomes, including asymptomatic colonization, mild diarrhea, and more severe disease syndromes such as pseudomembranous colitis, toxic megacolon, bowel perforation, sepsis, and death^3^. Antibiotics serve as the standard treatment for primary CDI^4,5^. However, CDI recurrence occurs in approximately a quarter of cases after antibiotic treatment^6,7^. Once CDI recurs, patients may get into a vicious cycle of antibiotic therapy and relapse^8^. Moreover, the use of antibiotics has been identified as the primary risk factor for developing CDI, and reports of strains with decreased sensitivity to vancomycin are becoming more frequent.

The human gut microbiome is critical in providing colonization resistance against exogenous pathogens through complex mechanisms such as nutrient competition, competitive metabolic interactions, niche exclusion, and induction of the host immune response^9^. Intestinal microbiota restoration, such as fecal microbiota transplantation (FMT), has been shown to be effective for CDI treatment as well as the restoration of colonization resistance against *C. difficile*^10,11^. While FMT has emerged as a promising therapy for recurrent CDI (rCDI), its exact mechanisms of action are not fully understood^12^. In addition, FMT has the potential to transmit undetected or emerging pathogens, which may result in hospitalization or even death^13,14^. Recently, the FDA has approved fecal microbiota products (e.g., Rebyota^15^ and Vowst^16^) for the prevention of rCDI in individuals 18 years of age and older, following antibiotic treatment for rCDI. Rebyota is a room temperature shelf stable suspension of healthy donor stool^17^, although its clinical effect size for the prevention of rCDI is modest (RR, 1.17; 95% CI, 0.99–1.39)^18^ and its microbial composition is not predefined^19^. Although Vowst is a formulation of live fecal microbiota consisting of a highly purified collection of about 50 species of *Firmicutes* spores with a more robust clinical effect size (1.46; 95% CI, 1.21– 1.75)^18^, the ecological principle underlying the selection of these microbial strains is unclear.

The variability of biological properties among bacterial strains within the same species underscores the significance of conducting strain-level composition analysis to understand the role of the human microbiome in human health and disease^20^. For example, some strains from *Escherichia coli* (e.g., *E. coli O157:H7*) cause severe abdominal pain, bloody diarrhea, and vomiting^21^. In contrast, *E. coli Nissle 1917* is a non-pathogenic strain that has been utilized as a probiotic agent to treat gastrointestinal infections in humans^22,23^. Whole metagenome shotgun (WMS) sequencing is a rapid, cost-effective, and high-throughput technology for profiling microbial communities in human microbiome studies^24^. However, precise identification of microorganisms at the strain level remains challenging. Additionally, traditional strain-level profilers can only identify strains within the reference genome databases^25^. These databases are subject to limitations and biases and are unable to characterize microbes that do not have high-quality reference genomes. To resolve these limitations, an alternative strategy for WMS data analysis involves reconstructing metagenome-assembled genomes (MAGs) through *de novo* assembly and binning, offering the advantage of recovering genomes for uncultured microorganisms absent from current reference databases^26^.

In this study, we leveraged a novel computational framework^27^ we previously developed to rationally design a bacterial consortium against CDI (**Fig. 1**). The metagenome assembly and binning strategies were applied to reconstruct microbial population genomes directly from the microbiome samples of two independent CDI-related cohorts as well as the healthy controls from the Human Microbiome Project (HMP). Specifically, we sought to identify known and unknown taxa at the strain level, quantify the degree of donor strain engraftment, and design a candidate bacterial consortium against CDI.

**Fig. 1.**
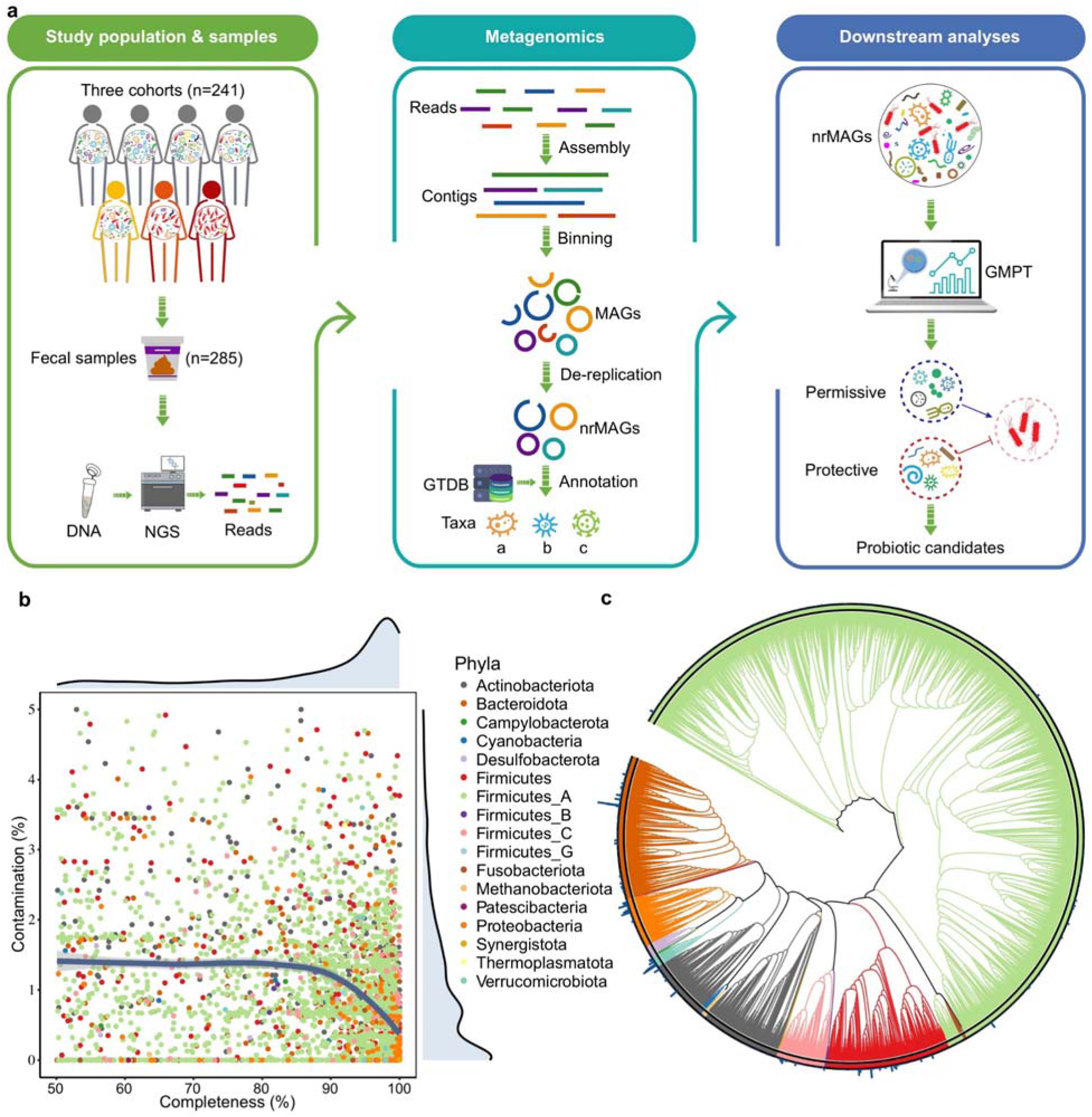
Study workflow and the reconstruction of the microbial genome catalog. **a.** To rationally design microbial consortia against *C. difficile*, we sought to infer species that may potentially inhibit *C. difficile* from various metagenomic data. We collected a total of 285 shotgun metagenomic sequencing data from three independent cohorts. A total of 7,769 MAGs (≥50% completeness and ≤5% contamination) were constructed from all metagenomic sequencing data. The MAGs were then dereplicated to 3,741 non-redundant MAGs (nrMAGs, strain level) based on 99% of ANI. The taxonomy annotation and abundance estimation of nrMAGs were then conducted. We then applied the generalized microbe-phenotype triangulation (GMPT) method to identify candidate strains for the development of microbiota probiotics. **b.** The distribution of completeness and contamination of nrMAGs is depicted, with the color of each point representing the respective phylum. Additionally, the size of each point corresponds to the genome size of the nrMAGs. **c.** A phylogenetic tree of nrMAGs was constructed using PhyloPhlAn. In this representation, the color of the outer cycle and clades signifies the phylum, while the bar plot within the cycle illustrates the average abundance across all microbiome samples.

## Results

### Study cohorts and metagenomic datasets

To rationally design microbial consortia against *C. difficile*, we aimed to infer species that may inhibit *C. difficile* from CDI-related microbiome samples. We first collected WMS sequencing data from our in-house clinical cohort (denoted as BIDMC-cohort hereafter) ^28,29^. Our BIDMC cohort consists of 104 well-characterized recruited participants divided into four groups (**Table S1** and **Fig. 2a**): (1) Control (CON, n = 26); (2) Non-CDI Diarrhea (NCD, n = 14); (3) Asymptomatic Carriage of *C. difficile* (ASC, n = 17); (4) CDI (n = 47). Given the fact that the participants from the CON group were not healthy people, we retrieved an FMT study^30^ (denoted as Verma-cohort hereafter) with publicly available data that assessed the microbiome composition of donors (n=21) and recipients (n=22, pre-and post-FMT) through WMS sequencing (**Methods** and **Fig. 2a**). In addition, we included two sets of randomly selected metagenome samples of healthy adults (n = 94) from the Human Microbiome Project (HMP)^31^.

**Fig. 2.**
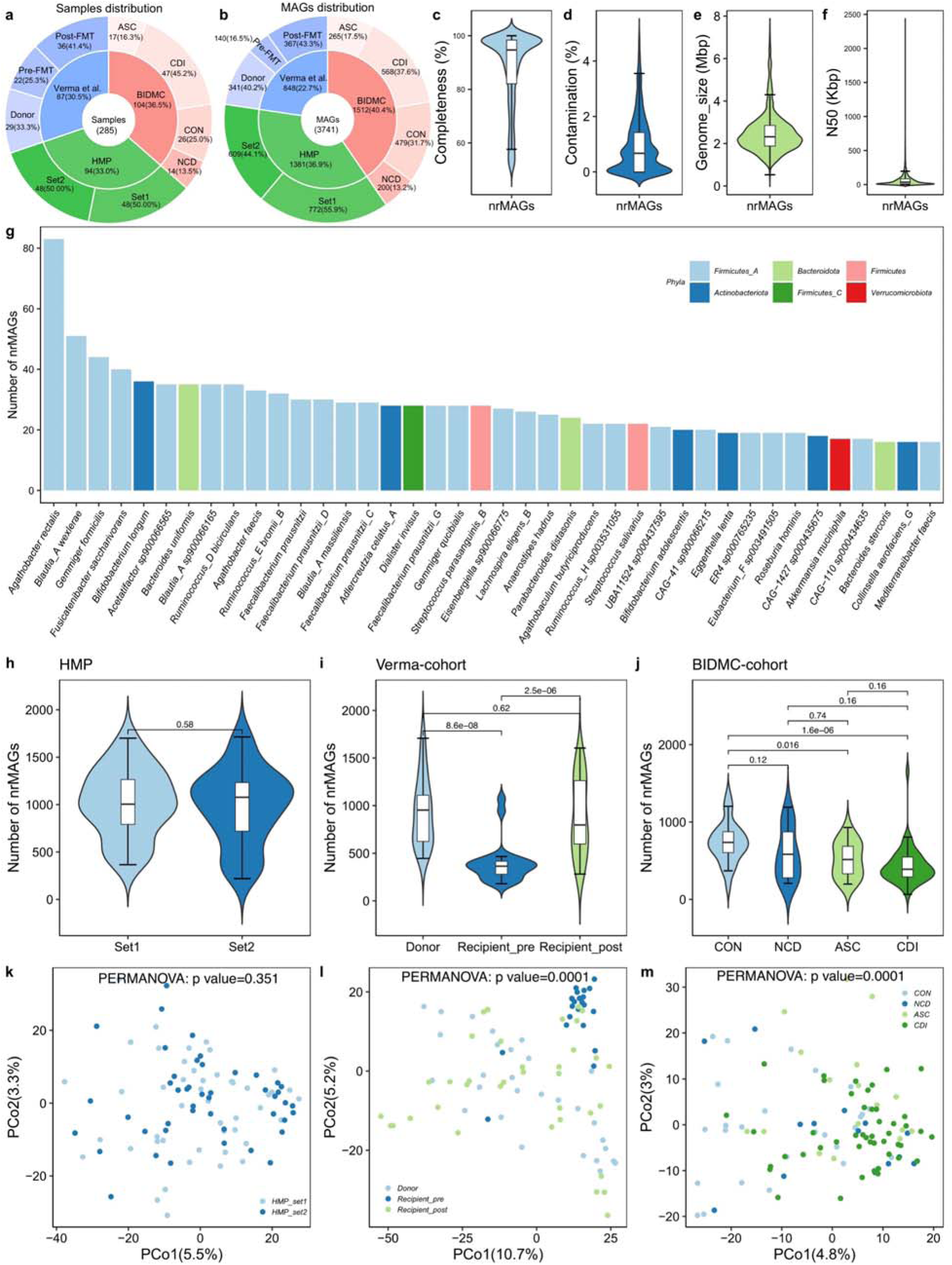
The microbial genome catalog and microbial diversity. **a.** Sample distribution among different datasets and clinical groups. **b**. Number of MAGs recovered from different datasets and clinical groups. Violin plot of basic characteristics of nrMAGs on completeness (**c**), contamination (**d**), genome size (**e**), and N50 (**f**). **g**. The top-40 species with the highest strain-richness (i.e., number of nrMAGs) identified from the microbial genome catalog. The color of each bar signifies the phylum. Richness (number of identified nrMAGs) of the gut microbiome from HMP (**h**), Verma-cohort. (**i**), and BIDMC-cohort (**j**). Principal Coordinates Analysis (PCoA) plot based on robust Aitchison distance from HMP (**k**), Verma-cohort. (**l**), and BIDMC-cohort (**m**). All PERMANOVA tests were performed with 9999 permutations based on robust Aitchison distance, two-sided.

### A high-quality microbial genome catalog

Following quality control, we performed metagenomic assembly and binning on those microbiome samples from three cohorts, yielding 7,769 MAGs. To evaluate the highest quality representative genomes, we dereplicated the 7,769 MAGs at an average nucleotide identity (ANI) threshold of 99%, resulting in a final set of 3,741 non-redundant MAGs (nrMAGs) with strain-level resolution. The nrMAGs were contributed by HMP, Verma-cohort, and the BIDMC-cohort in proportions of approximately 37%, 23%, and 40%, respectively **(Fig. 2b**). In particular, our findings indicate that recipients prior to FMT made a smaller contribution to the nrMAG collection compared to donors and recipients after FMT, suggesting a reduced microbial diversity **(Fig. S1**). These nrMAGs exhibited a mean completeness of 88%, mean contamination of 0.93%, mean genome size of 2.5 megabases (Mb), and mean N50 of 65.7 kilobases (kb) (**Fig. 1b-c** and **Fig. 2c-f**). Out of the 3,741 strain-level nrMAGs, 1,390 (37.16%) nrMAGs met medium-quality criteria (50% ≤ completeness < 90%, and ≤5% contamination), while 2,351 (62.84%) nrMAGs exhibited high-quality ( ≥ 90% completeness, and ≤ 5% contamination)^32,33^ (**Fig. 1b**).

Using the Genome Taxonomy Database^34^, these nrMAGs were taxonomically assigned to 17 phyla, 22 classes, 47 orders, 104 families, and 408 genera, spanning across 883 species. Most of them belonged to Firmicutes_A (60.97%), followed by Bacteroidetes (10.18%), and Actinobacteria (10.00%). The phyla information of nrMAGs was summarized in **Fig. 1b-c**. Among those 883 species, *Agathobacter rectalis*, *Blautia_A wexlerae*, *Gemmiger formicilis*, *Fusicatenibacter saccharivorans*, and *Bifidobacterium longum* were the top five species with the highest strain-level diversity (i.e., number of nrMAGs identified within a specific species, **Fig. 2g**).

### Microbial diversity

We initially investigated alpha diversity in the human microbiome at the nrMAG level. Alpha diversity measures (i.e., Richness and Shannon index) were compared among different groups from the same cohorts (**Fig. 2h-j**). No significant differences were found between the two randomly selected sets from the HMP (**Fig. 2h** and **Fig. S2a**). In accordance with the original study^30^, we found that the Richness and Shannon indices of the gut microbiome in the recipients of pre-FMT were significantly lower than those in donors. After FMT, those recipients showed similar alpha diversity to donors (**Fig. 2i** and **Fig. S2b**). In the BIDMC-cohort, we found that only the CDI group showed significantly lower alpha diversity than the CON group (**Fig. 2j** and **Fig. S2c**). Participants from the ASC group only showed a significantly lower number of identified nrMAGs than CON group participants.

Principal coordinate analysis (PCoA) based on robust Aitchison distance, combined with PERMANOVA (permutational multivariate analysis of variance, a statistical method commonly used for testing the association between the microbiome and a covariate of interest), revealed no significant difference in the gut microbial community structure at the nrMAG-level between the two datasets from HMP (**Fig. 2k**). We found that the microbiomes of the donor and the recipients from pre-and post-FMT were compositionally distinct in the Verma-cohort (*P* = 0.0001, PERMANOVA **Fig. 2l**). Consistent with our previous study using 16S rRNA gene sequencing data^29^, the overall microbial composition differed significantly among different groups in the BIDMC-cohort (*P* = 0.0001, PERMANOVA **Fig. 2m**).

### Identify potential permissive and protective nrMAGs for the rational design of live biotherapeutics

To identify candidate strains for the development of microbiota-derived biotherapeutics, we applied the generalized microbe-phenotype triangulation (GMPT) method, moving beyond the standard association analysis^27^. The GMPT relies on the following core hypothesis: species that are differentially abundant in most pairwise phenotype-based comparisons and whose abundances display a strong negative (or positive) correlation with the abundance of the pathogen tend to be causal preventive (or permissive) species that directly inhibit (or promote) the growth of the pathogen^27^. Our GMPT analysis incorporated microbiome data from the BIDMC-cohort, donor data from the Verma-cohort^30^, and one set from HMP. Since we have two sets of randomly selected metagenome samples of healthy adults from the HMP, we systematically included one set of HMP microbiome data at a time to cross-validate the results between the two datasets. Then, we conducted pairwise comparisons for six individual phenotype groups, which encompassed CDI, ASC, NCD, and CON from the BIDMC-cohort, donors from the Verma-cohort, and a dataset from HMP.

Applying this approach to the data with the first set of HMP data, 15 pairwise differential abundance analyses generated a total of 1,349 nrMAGs present in at least one pairwise comparison (**Table S6**). To explore the potential relationship between those candidate nrMAGs and CDI, we calculated Spearman correlation coefficients between the average relative abundances of nrMAGs and pragmatic severity scores in a continuum of non-CDI controls and *C. difficile* colonized and infected subjects (i.e., HMP healthy controls: 0; Donor from Verma et al.^30^: 1; CON: 2; NCD: 3; ASC: 4 and CDI: 5) in different phenotypes. Similarly, we identified a total of 1,390 nrMAGs present in at least one pairwise comparison with the second set of HMP data (**Table S7**). Among the protective nrMAGs between the two runs with HMP data, 80.77% (525/650) and 81.14% (525/647) of them were overlapped, respectively. We then computed the average rank between two runs based on the frequency (**Table S8)**. Among the top 40 potential protective nrMAGs, the dominant species were *Dorea formicigenerans*, *Oscillibacter welbionis*, *Faecalibacterium prausnitzii*, *GCA-900066135 sp900066135*, *Bariatricus comes*, *Phocaeicola dorei*, *Anaerobutyricum hallii*, *Bacteroides ovatus*, *Blautia_A obeum*, *Mediterraneibacter faecis*, *Alistipes putredinis*, *Odoribacter splanchnicus*, *Streptococcus salivarius*, and *Dorea longicatena* (**Table 1**). Through a systematic review of literature, we found that most of our candidate strains have been reported to be protective from CDI or non-CDI antibiotic associated diarrhea at higher taxonomical levels (e.g., species and genus levels) across existing studies (**Table 1**). These findings support the validity of our methods.

**Table. 1.**
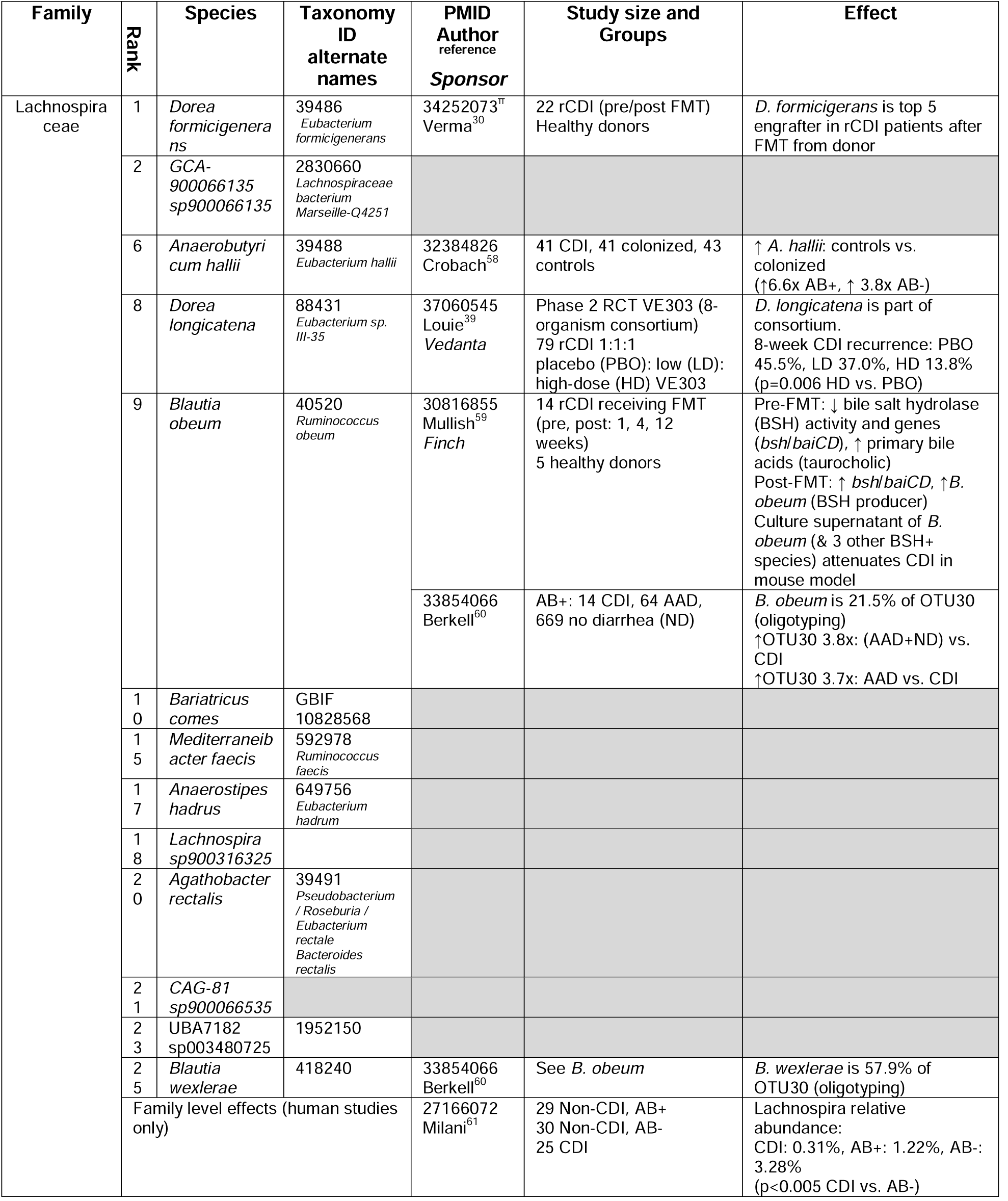
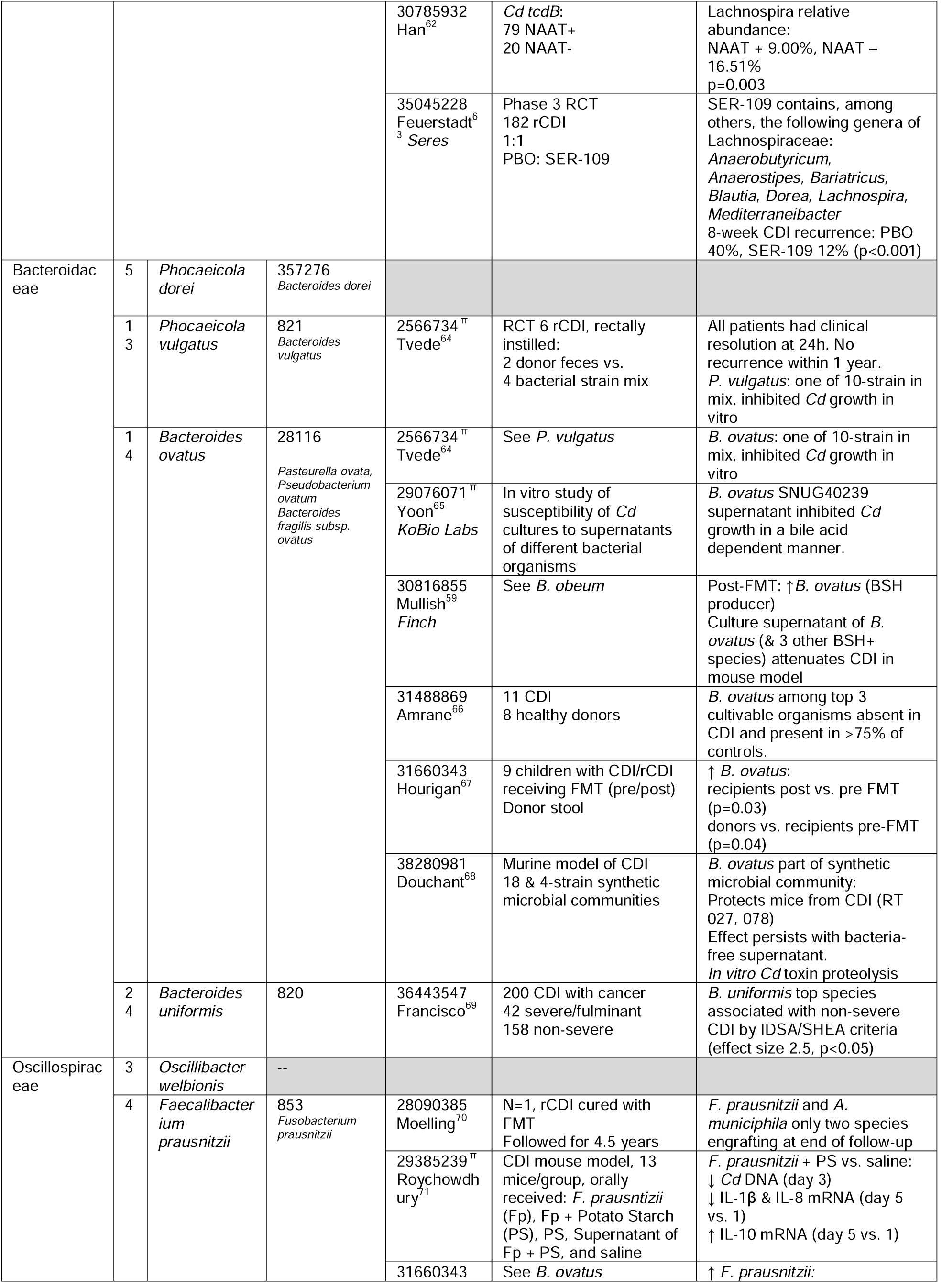
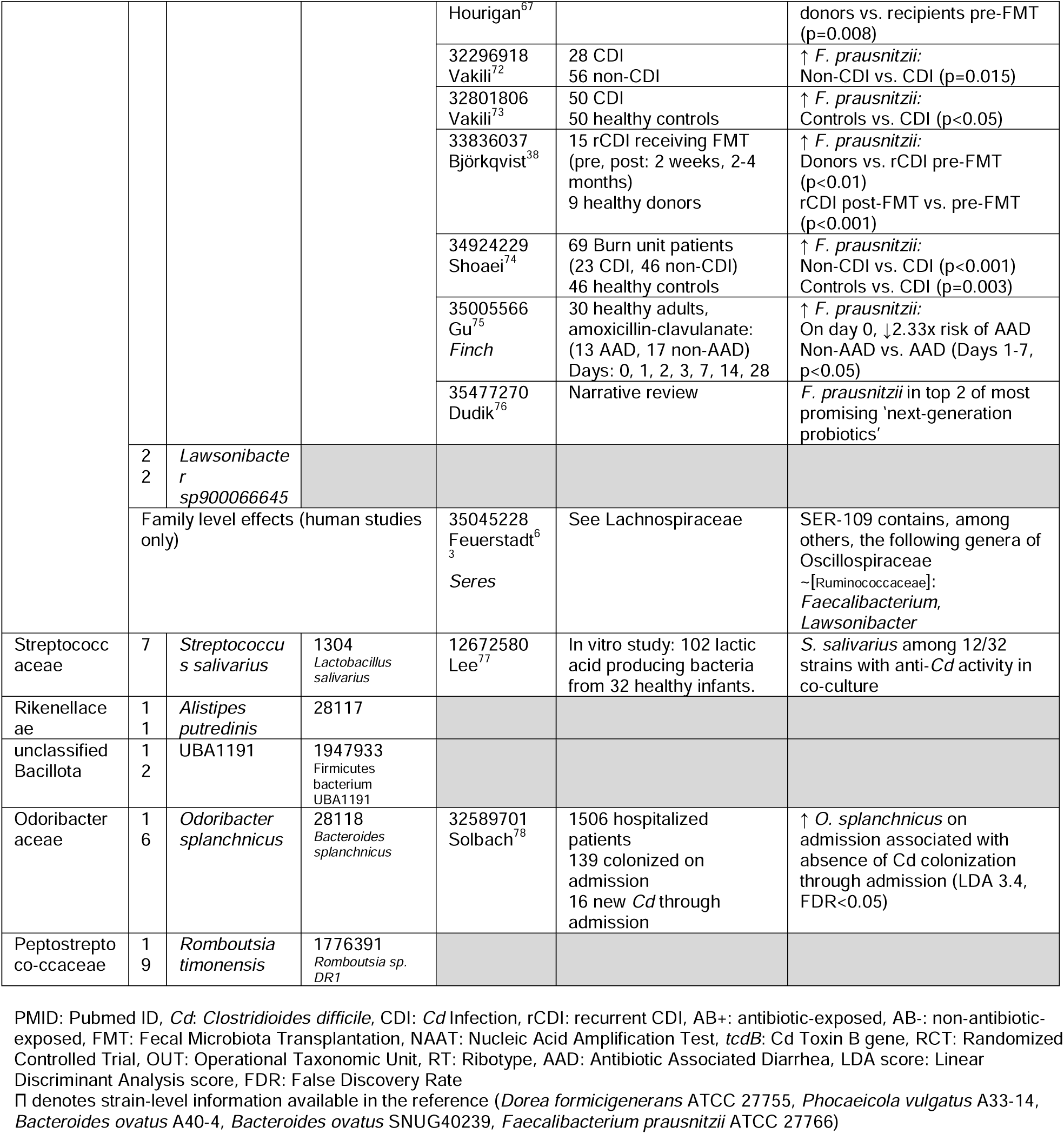
Summary of the literature evidence regarding the potential role of protective species in CDI identified through our computational pipeline. The top 25 potential protective species were selected based on the overlapped protective strains identified from the GMPT results with two sets of HMP microbiome data.

### The protective strains play an important role in FMT

To further validate the potential role of the protective strains we identified from the GMPT pipeline, we systemically tracked the microbiome changes of the recipients who underwent FMT in the Verma-cohort. We aimed to investigate if those protective strains also play an important role in the success of FMT. Notably, the microbiome samples from the recipients in the Verma cohort were not included in our previous GMPT analysis.

First, we examined the gain and loss of microbial strains before and after FMT to assess the transfer and engraftment of the donor microbiome in the recipient. For donor, pre-FMT recipients, and post-FMT recipients, we identified 3,129, 2,093, and 3,054 nrMAGs, respectively. Notably, post-FMT recipients showed a loss of 33 nrMAGs (**Fig. 3a**), with the majority of the lost strains attributed to species such as *Anaeroglobus micronuciformis*, *Phascolarctobacterium faecium*, *Fusobacterium polymorphum*, and *Duodenibacillus sp003472385* from Firmicutes_C, Proteobacteria, and Fusobacteriota (**Fig. 3b**). On the contrary, all recipients exhibited a gain of 923 nrMAGs from their donors (**Fig. 3a**). The majority of these engrafted strains were taxonomically annotated to Actinobacteriota and Firmicutes_A, such as species like *Ruminococcus_D bicirculans*, *Faecalibacterium prausnitzii*, *Faecalibacterium prausnitzii_G*, *Agathobacter rectalis*, *Agathobacter faecis*, *Acetatifactor sp900066565*, *Bifidobacterium adolescentis*, and *Collinsella aerofaciens_G* (**Fig. 3c-d**).

**Fig. 3.**
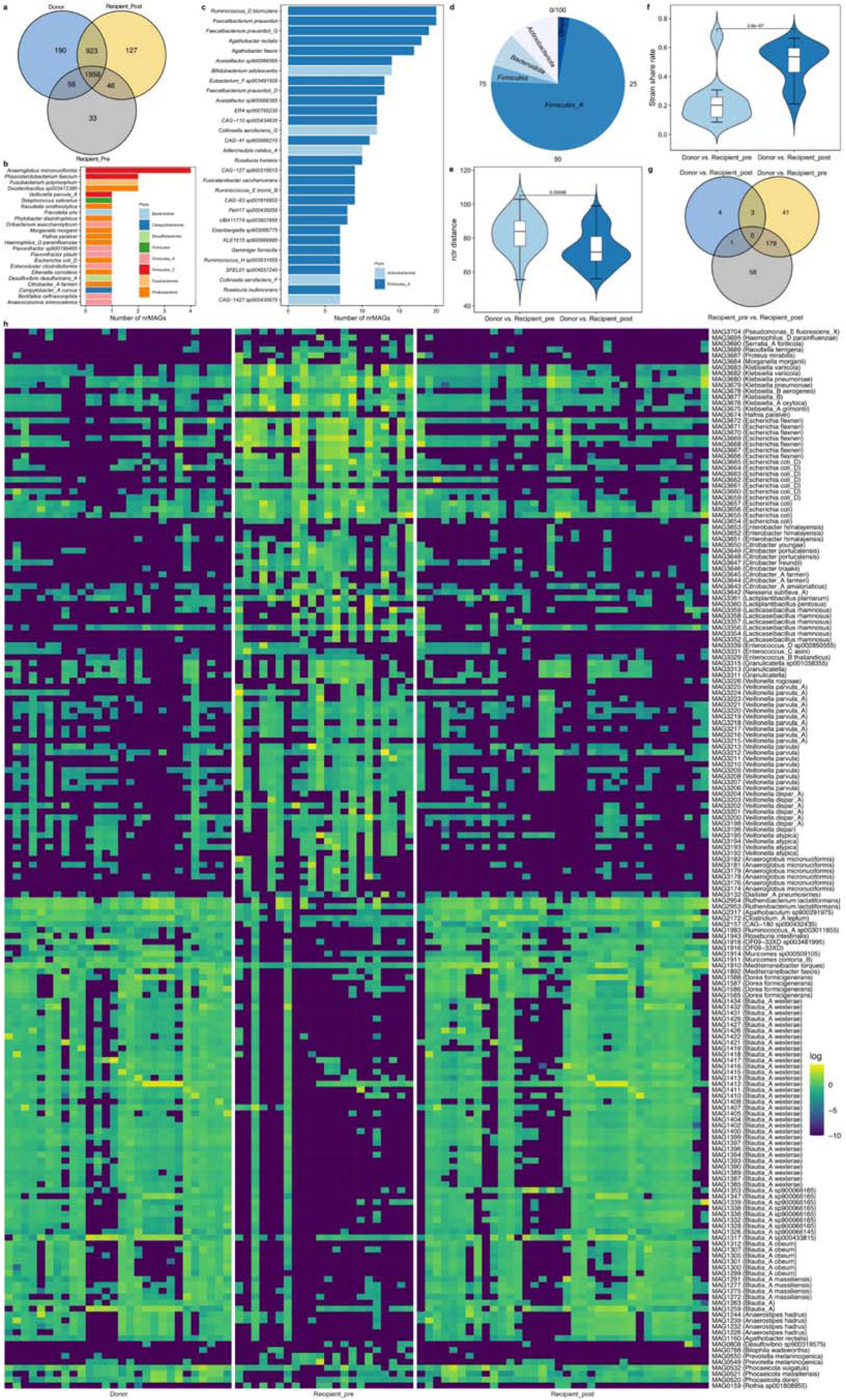
The changes in recipients’ microbiome after FMT. **a**. The distribution of nrMAG among donors, pre-FMT recipients, and post-FMT recipients. **b**, The distribution of lost nrMAGs after FMT at the species level, and the color of each bar represents the phylum. **c**. The distribution of engrafted nrMAGs after FMT at the species level, and the color of each bar represents the phylum. **d**. The distribution of engrafted nrMAGs after FMT at the phylum level. **e**, The robust Aitchison distance between donor and recipient pairs before and after FMT. **f**, The nrMAG share rate between donor and recipient pairs before and after FMT. **g**. The differential abundant nrMAG distribution among three pair-wise comparisons between donors, pre-FMT recipients, and post-FMT recipients. **h**. The heat map showed the abundance distribution of overlapped nrMAGs identified from the comparisons of donor vs. pre-FMT recipients and pre-FMT recipients vs. post-FMT recipients. These nrMAGs were taxonomically annotated using GTDB-Tk based on the Genome Taxonomy Database.

For each donor-recipient pair, we then calculated the difference in their gut microbial community structure before and after FMT using the robust Aitchison distance. Our findings indicate that the distance between donors and recipients was significantly reduced after FMT compared to the pre-FMT state (**Fig. 3e**). Additionally, calculating the strain share rate for each donor-recipient pair before and after FMT revealed agreement with our previous finding that recipients gained more strains, shared a greater number of strains with the donor after FMT (**Fig. 3f**).

Changes in the microbiome induced by FMT not only indicate the transfer and engraftment of the donor microbiome but also involve alterations in the abundance of coexisting strains. To address this question, we conducted the differential abundance analysis among three groups. Consistent with the robust Aitchison distance and strain share rate analyses, we only identified less differential abundant strains between donor and post-FMT recipients (**Fig. 3g and Table S2**). We have identified 223 and 238 differential abundant nrMAGs from the comparison of donor vs. pre-FMT recipients (**Fig. 3g and Table S3**) and pre-FMT recipients vs. post-FMT recipients (**Fig. 3g and Table S4**), respectively. Among these differential abundant nrMAGs, we found 179 overlapped strains (**Fig. 3h, Table S5**), including strains from *Blautia_A wexlerae*, *Veillonella parvula_A*, *Veillonella parvula*, *Blautia_A sp900066165*, *Escherichia coli_D*, *Escherichia flexneri*, *Anaeroglobus micronuciformis*, *Blautia_A obeum*, *Lacticaseibacillus rhamnosus*, and *Veillonella dispar_A.* Specifically, we observed significant increases in some candidate protective strains following FMT. These include multiple strains from *Dorea formicigenerans*, *Mediterraneibacter faecis*, *Phocaeicola dorei*, *Blautia_A wexlerae*, and *Blautia_A obeum*. This finding further validated the potential role of protective strains in treating CDI.

## Discussion

The growing interest in FMT as a therapeutic approach stems from its high success rate in treating recurrent CDI, leading to an exploration of its potential for addressing various human diseases^35^. However, FMT remains an unstandardized procedure with unclear mechanisms and long-term safety concerns^35,36^. Therefore, an advantage of microbial consortia over “whole stool” FMT is the introduction of a group of specific microbiota that can precisely target and effectively treat a disease while minimizing clinical risks. In this study, we used a computational pipeline to directly identify candidate bacterial strains from a diverse CDI-related metagenomic dataset, thereby facilitating the targeted development of microbial therapies and advancing our understanding of CDI pathogenesis and treatment.

By tracking the dynamic changes in gut microbiome data undergoing FMT, we identified significant shifts in the microbial structure of the recipients. Although we did not utilize the microbiome data from recipients before and after FMT in our GMPT pipeline, we found that some of the top ranked candidate protective strains showed significant increases after FMT, including multiple strains from *Dorea formicigenerans*, *Mediterraneibacter faecis*, *Phocaeicola dorei*, and *Blautia_A obeum*. This finding provides an additional layer of validity to our method. In addition, we performed a systematic literature review on the highest ranked candidate protective strains and found that the majority of them have been reported to have various protective roles at species or genus levels in the CDI continuum: negative association with *C. difficile* colonization, infection and severity. We found clustering of the main protective species within the families Lachnospiraceae, Bacteroidaceae, and Oscillospiraceae. For example, *F. prausnitzii*, a beneficial human gut microbe touted as a candidate for next-generation probiotics^37^, was found to have reduced abundance in CDI patients, which was restored after FMT^38^. Interestingly, we have also identified a protective strain of the species *Dorea longicatena,* which is a component of a defined bacterial consortium (VE303) with encouraging Phase 2 clinical data, consisting of eight, nonpathogenic, nontoxigenic, commensal strains of Clostridia^39^.

In addition to the potential protective strains, we also identified multiple permissive strains of *C. difficile*, including strains from *Enterococcus_B faecium* and *Eggerthella lenta*. This aligns with a previous study reporting that enterococci (including *E. faecium*) can enhance the fitness and pathogenicity of *C. difficile* via shaping the metabolic environment in the gut and reprograming *C. difficile* metabolism^40^. Additionally, *E. lenta* is an anaerobic gram-positive bacillus associated with polymicrobial intraabdominal infections^41^. Therefore, the potentially permissive strains that we identified from this study offer the opportunity to further understand how *C. difficile* interacts with the rich community of microorganisms in the colon. Moreover, in alignment with the variation in biological properties among bacterial strains within the same species^42^, we have observed distinct roles played by different strains of *F. prausnitzii_D* in the context of CDI. This underscores the critical importance of conducting studies at the strain level.

The current study has some limitations. First, we leveraged metagenomic data from three independent datasets with technological variations, including differences in sequencing depth. Second, we did not pre-define a strict threshold to select potential protective strains from the candidate list for further experimental validations. Lastly, the inference of the efficacy of candidate protective strains against CDI is limited by the current computational algorithm. To test the efficacy of our proposed microbial consortia and gain a deeper understanding of exact mechanisms, the utilization of techniques of metabolomics and immunological approaches, along with direct *in vitro* and *in vivo* experiments, are necessary.

Taken together, our results provide compelling evidence for the rational design of microbial consortia against *C. difficile*. Many of the candidates detected here replicate previously reported findings, supporting the validity of our results. Importantly, our work paves the way for the design of LBPs against general microbiome-related diseases.

## Methods

### Study cohorts

#### Dataset I: BIDMC-cohort

The background and design of this cohort have been detailed in our previous studies^28,29^. This clinical cohort consists of 104 well-characterized recruited participants, who were divided into four groups associated with different *C. difficile* infection/colonization statuses: (1) *C. difficile* infection (CDI, n=47): Eligible patients were inpatients ≥ 18 years old with new-onset diarrhea, positive clinical stool NAAT (Xpert *C. difficile*/Epi) result, and a decision to treat for CDI; (2) Asymptomatic Carriage (ASC, n = 17): Eligible patients were inpatients ≥ 18 years old, admitted for at least 72 hours, who had received at least one dose of an antibiotic within the past seven days, and did not have diarrhea in the 48 hours prior to stool specimen submission, and positive clinical stool NAAT result; (3) Non-CDI Diarrhea (NCD, n = 14): patients with diarrhea (confirmed using the same definition used for the CDI cohort) but had NAAT-negative stool on clinical C. difficile testing; and (4) Control (CON, n = 26): patients without diarrhea who had screened as eligible for the ASC cohort but were NAAT-negative on research stool testing. DNA of fecal samples (200 mg) were extracted using Mag-Bind® Universal Metagenomics Kit (Product# M5633-01, Omega Biotek) and DNeasy PowerSoil Kit (Catalog# 12888-100, Qiagen) according to manufacturer’s instructions. The quality of the extracted DNA was measured by 1% agarose gel electrophoresis and Qubit® 3.0 Fluorometer (ThermoFisher). Subsequently, the extracted DNAs were used for shotgun metagenomic library construction, and sequencing was performed on the Illumina HiSeq X Ten platform, generating a 150 bp paired-end library for each sample.

#### Dataset II: Verma-cohort

In the study conducted by Verma et al^30^, fecal samples were collected from 22 patients with recurrent CDI before and after FMT and their corresponding healthy donors (n=21, with one donor providing fecal samples for two different recipients). Eight-seven WMS human gut metagenomes were downloaded from this study via NCBI Sequence Read Archive (BioProject ID PRJNA705895). The clinical outcome in recurrent CDI patients after FMT was determined by the symptomatic resolution of CDI^30^. Clinical symptoms such as diarrhea, bloating, abdominal pain, and cramping were alleviated in all patients within 3–7 days following FMT^30^.

#### Dataset III

Human Microbiome Project. Human gut metagenomes (Ninety-eight individuals) were randomly selected from HMP data (https://portal.hmpdacc.org/)^43^. All samples are from the HMP study^31^ and are healthy adult subjects. In total, ninety-four human gut metagenomes were randomly selected based on the largest group size in our clinical cohort. To cross-validate the main findings, we randomly divided the HMP data into two sets in the downstream analyses.

### Metagenome assembly and binning

Genome reconstruction of the human microbiome using metagenomic sequencing data was executed through the functional modules of metaWRAP (v1.3.2)^44^. All metagenomic sequencing data underwent quality control and removal of human contamination using metaWRAP-Read_qc. Clean reads were then assembled with the metaWRAP-Assembly module using metaSPAdes (v3.13.0)^45^. The assembled contigs were binned into bins using three metagenomic binning tools: MetaBAT (v2.12.1)^46^, MaxBin (v2.2.6)^47^, and CONCOCT (v1.0.0)^48^. The default minimum length of contigs used for constructing bins with MaxBin2 and CONCOCT was 1000 bp, and metaBAT2 was defaulted to 1500 bp^44^. The bins from each binning tool were integrated and refined with Bin_refinement module of metaWRAP with options “-c 50 -x 10”, corresponding to the criterion of medium-quality draft MAGs^32^. CheckM (v1.0.12)^49^ was used to estimate the completeness and contamination of the bins, and the minimum completion and maximum contamination were 50% and 10%, respectively.

### De-replication of MAGs and genome annotation

All 7,776 MAGs underwent de-replication into non-redundant MAGs (nrMAGs) using dRep (v3.0.0) (≥50% genome completeness and ≤5% contamination)^50^. Initially, MAGs from three cohorts were divided into primary clusters using Mash at a 90% Mash ANI. Then, each primary cluster was used to form secondary clusters at the threshold of 99% ANI with at least 30% overlap between genomes^51^. Taxonomic annotation of all nrMAGs was conducted using GTDB-Tk (v.1.4.1)^52^ based on the Genome Taxonomy Database (http://gtdb.ecogenomic.org/)^34^, providing standardized taxonomic labels for subsequent analysis in this study.

### Abundance estimation and phylogenetic analysis of nrMAGs

The metaWRAP-Quant_bins module coupled with Salmon (v0.13.1)^53^ was employed to access the abundance of each nrMAGs in each metagenomic sample. The phylogenetic tree of the nrMAGs was constructed using PhyloPhlAn (v3.0.58)^54^ and visualized through iTOL (https://itol.embl.de/)^55^.

### Statistical analysis

Microbial alpha diversity measures were calculated at the nrMAGs level using R vegan package (v2.5.7), and principal coordinates analysis (PCoA) plots were generated using robust Aitchison distance^56^. Differences in microbiome compositions across different groups were tested by the permutational multivariate analysis of variance (PERMANOVA) using the “adonis” function in R vegan package. All PERMANOVA tests were performed with 9999 permutations based on the robust Aitchison distance. We defined strain-sharing rates as the total number of shared strains between two samples divided by the number of common species identified from the two samples. Differences between the groups were analyzed using a Wilcoxon–Mann–Whitney test. For differential abundance analysis and GMPT (Generalized Microbe Phenotype Triangulation) pipeline^27^, we used ANCOM (analysis of composition of microbiomes)^57^, with a Benjamini–Hochberg correction at a 5% level of significance. All statistical analysis was performed with R (version 3.6.3).

## Data availability

Metagenomic data from HMP are available via https://portal.hmpdacc.org. The metagenomic data from the study of Verma et al.^30^ can be downloaded via NCBI Sequence Read Archive (BioProject ID PRJNA705895). The metagenomic data from the BIDMC-cohort is available in the NCBI Bioproject under accession code PRJNA1067975. Metagenome-assembled genomes for all samples are available on Figshare (https://doi.org/10.6084/m9.figshare.25355857).

## Code availability

The code for the construction of the MAGs catalog and statistical analysis and visualization is available in the GitHub repository (https://github.com/ShanlinKe/CDI).

## Acknowledgements

The authors thank all patients who participated in this study, as well as the technologists in the Beth Israel Deaconess Medical Center Clinical Microbiology Laboratory, for their help with sample collection. This study was supported by the China Scholarship Council (201806305024 to Y.C.), National Key Research and Development Program (grants 2021YFD1300301 to Y.C.), Shaanxi Science Fund for Distinguished Young Scholars (grants 2024-JC-JCQN-25 to Y.C.), the Ningxia Key Project of Research and Development Plan (grants 2023BFC01036 Y.C.), the National Institute of Allergy and Infectious Diseases (grants R01AI141529 to Y.-Y.L., R01AI116596 to N.R.P. and C.P.K., and K23AI177749-01 to J.A.V.G.), Institute Merieux (grant to N. R. P. and C. P. K.), the Irving W. and Charlotte F. Rabb Award (to X.C.) and Milky Way Life Sciences (contract to C.P.K. and Y.-Y.L).

## Author contributions

Y.-Y.L., X.C. and C.P.K. conceived and designed the project. N.R.P. and C.P.K. planned and performed the human studies and sample collections. S.K. performed all the data analysis. S.K., J.A.V.G., and Y.-Y.L. interpreted the results and prepared the manuscript. Z.S., X.C., and C.P.K. interpreted the results, reviewed and edited the manuscript. Y.C. acquired the raw sequencing data, reviewed and edited the manuscript. All authors have read and approved the manuscript.

